# Distinction between small RNA-bound and free ARGONAUTE via an N-terminal protein-protein interaction site

**DOI:** 10.1101/2022.10.22.513346

**Authors:** Simon Bressendorff, Ida Marie Zobbe Sjøgaard, Andreas Prestel, Birthe B. Kragelund, Christian Poulsen, Peter Brodersen

## Abstract

ARGONAUTE (AGO) proteins bind to small non-coding RNAs to form RNA Induced Silencing Complexes (RISCs). In the RNA-bound state, AGO proteins are stable while RNA-free AGOs turn over rapidly. Molecular determinants unique to RNA-free AGO that allow its specific recognition and degradation remain unknown. Here, we show that a confined, linear region in Arabidopsis AGO1, the N-coil, is accessible to antibodies preferentially in the RNA-free state of AGO1. Reanalysis of hydrogen-deuterium exchange data on human Ago2 indicates similar structural flexibility of the N-coil depending on small RNA binding. Unloaded Arabidopsis AGO1 interacts with the autophagy cargo receptor ATI1 via direct contact to specific amino acid residues in the N-coil, and mutation of residues required for ATI1 interaction reduces the degradation rate of unloaded AGO1 *in vivo*. These results provide insight into the molecular basis for specific recognition and degradation of the RNA-free state of eukaryotic AGO proteins.

## INTRODUCTION

RNA Induced Silencing Complexes (RISCs) are central components in the processing of genetic information in eukaryotes. The core of RISC is an ARGONAUTE (AGO) protein bound to a small non-coding RNA that guides RISC to complementary target RNAs by base pairing. In turn, RISCs repress gene expression sequence-specifically using a variety of mechanisms operating at either transcriptional or post-transcriptional levels. In animals, two distinct phylogenetic clades of AGO proteins exist. The Ago clade carries out endogenous gene regulation, in the main via microRNAs (miRNAs)^1^, and, in particular in invertebrates, basal antiviral defense via virus-derived small interfering RNAs (siRNAs)^2^. The Piwi clade is primarily used for silencing of transposable elements in the germline via Piwi-interacting RNAs (piRNAs)^3^. Plants do not encode Piwi-clade proteins, and use Ago-clade proteins to effect transposon silencing, basal antiviral defense, and endogenous gene regulation via miRNAs and siRNAs^4^. AGO1, the founding member of the AGO protein family, has widespread functions in small RNA-guided regulation at the post-transcriptional level, and is implicated in endogenous gene regulation via most miRNAs and in antiviral defense via virus-derived siRNAs^5^.

All AGO proteins consist of four separate domains called N, PAZ, MID and Piwi^6^. They fold into a two-lobed structure in the small RNA-bound state with N and PAZ together connected by the linker region L1 in one lobe and MID/Piwi in the other^7-10^ (Figure 1A,B). The ends of the small RNA are tethered in dedicated binding pockets in the MID and PAZ domains such that the 5’-end is bound to MID^11,12^ and the 3’-end to PAZ^13^. PAZ and MID are connected by a long linker sequence called L2 that also interacts extensively with the small RNA backbone and with other AGO domains in the small RNA-bound state ^8-10^ (Figure 1A,B).

**Figure 1.**
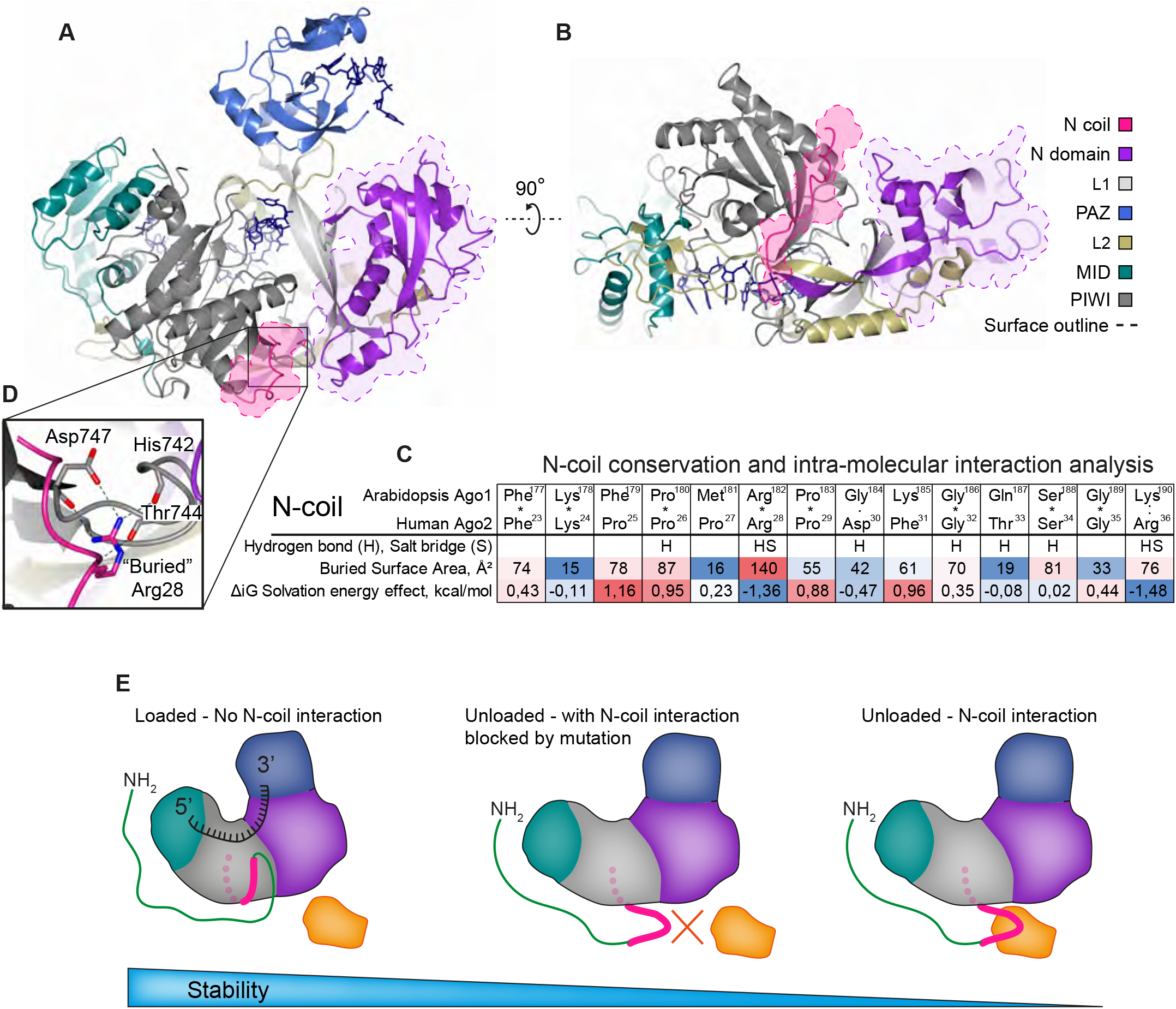
Structural properties and conserved interactions of the N-coil in Hs-Ago2. (A)-(B) Overview of the *Hs*Ago2 domain structure with surface outline of the globular N domain and the N-coil. (A) “side view”; (B) “bottom view”. (C) Intramolecular interactions between the N-coil and the remaining part of *Hs*Ago2 as calculated by the Protein Interfaces, Surfaces and Assemblies service PISA^59^. The solvation energy gain of the N-coil is calculated as the difference between free energy of the Piwi-associated and -dissociated N-coil. A negative solvation energy, Δ_i_G, of a residue makes a positive contribution to the solvation energy gain of the interface, thus favoring the dissociated state. The color gradient (blue to red) indicates the effect sizes of buried surface area and solvation energy and highlights the unique properties of Arg-28 in *Hs*Ago2. (D) Inlet showing conserved residues and bonds between a buried arginine side chain in the N-coil (Arg-28 in Hs-Ago2) and nearby residues. (E) Model illustrating a hypothetical function of the N-coil in distinguishing loaded from unloaded AGO protein and in conferring rapid proteolysis on unloaded AGO. Left, in the loaded form, the N-coil is inaccessible for protein-protein interaction due to its association with the Piwi domain. In the unloaded form (middle and right), the N-coil is accessible for interactors, leading to rapid proteolysis (right) that can be slowed down if N-coil-dependent interactions are precluded by mutation (middle). Presentations and PISA calculations were made using the crystal structure coordinates of the miR20a-bound *Hs*Ago2 (PDB 4F3T)^10^.

The chaperone-assisted assembly of RISC by incorporation of a small RNA into AGO, referred to as “RISC loading”, is a critical event in all RNA silencing pathways. It is rate-limiting for maturation of many miRNAs *in vivo*^14^, and in both metazoans and plants, it is decisive for the stability of AGO proteins. Small RNA-bound AGO is stable, but unloaded AGO is rapidly degraded *in vivo*^15-17^, unless it is bound to Hsp90^17,18^. Both the ubiquitin-proteasome pathway and autophagy are involved in turnover of unloaded AGO in plants and animals^15-20^, and progress has been made on identifying molecular components mediating such degradation. For example, the RING finger protein Iruka ubiquitinates the fly miRNA effector Ago1 in the L2 region specifically in its unloaded form ^21^. This ubiquitination triggers binding of an adaptor complex, ultimately leading to association with the valosin-containing protein (VCP) that mediates degradation of the ubiquitinated, unloaded Ago1 via autophagy^20^. A clear identification of sites differentially accessible between loaded and unloaded Ago1 was not achieved, however, and it remains possible that Iruka recognizes unloaded Ago1 outside of the ubiquitinated L2 region. Moreover, the Ago1-Iruka case is unlikely to be easily generalized to other systems, because the site of ubiquitination is non-conserved, and is situated in a *Dm*Ago1-specific insertion in L2 compared to other miRNA-interacting AGO proteins (Figure S1). In arabidopsis, mutation of the F-box protein FBW2 restores steady-state levels of a largely unloaded mutant AGO1 protein with a destabilized fold of the PAZ domain^22^. It is, however, unclear whether FBW2 targets AGO1 directly, and whether it specifically mediates degradation of unloaded, but functional AGO1, or plays a more general role in chaperone-assisted protein degradation elicited as a consequence of partial unfolding of the PAZ domain in the point mutant analyzed. Thus, despite considerable progress on identifying the downstream pathways mediating degradation of unloaded Ago proteins, the molecular determinants of unloaded AGO that distinguish it from loaded AGO and allow its selective recognition remain unknown.

In this study, we use arabidopsis AGO1 to define a region of crucial importance in the distinction between eukaryotic AGO proteins in the RNA-bound and in the free state. We show that the N-coil, a small, highly conserved linear region in the N domain, yet part of the MID-Piwi lobe in the loaded conformation, is preferentially accessible in unloaded arabidopsis AGO1 and human Ago2. The N-coil in arabidopsis AGO1 functions as a direct binding site for the autophagy cargo receptor ATG8 INTERACTING PROTEIN1 (ATI1), and mutations disrupting ATI1 binding reduce the degradation rate of unloaded AGO1 *in vivo*. These results highlight the N-coil as a key determinant of the distinction between loaded and unloaded AGO1, and, given its conserved structural properties, probably in other eukaryotic AGO proteins in both Ago and Piwi clades.

## RESULTS

### Definition of the N-coil and other N-terminal parts of arabidopsis AGO1

*Arabidopsis thaliana* (At) AGO1 has a 176-aa extension at its N-terminus, in addition to the canonical N-PAZ-MID-PIWI domains found in all AGO proteins. Because this extension is predicted to be an intrinsically disordered region (IDR, Figure S2), we term this part of AGO1 the N-IDR. The N-IDR is followed by a 14-aa region (Phe177-Lys190) that we previously termed the N-coil^5^. Structurally, the N-coil is of interest, because it is part of the MID-PIWI lobe in AGO-small RNA complexes, despite its location in the N-terminal part of AGO in sequence. (Figure 1A-B). Furthermore, the N-coil assumes an extended conformation with no secondary structure, yet has low structural flexibility, and reaches across the width of the entire AGO-small RNA complex (Figure 1B). The N-coil is followed by two *β*-strands that connect it to the remainder of the N-domain in the N-PAZ lobe (Figure 1B). We term this latter part of the N-domain the globular N-domain to emphasize its globular fold, in contrast to the N-coil.

### Properties of the N-coil and a hypothesis for its function

Two additional properties of the N-coil make this part of AGO proteins attractive for further study. First, residues in the conserved N-coil are structurally fixed by numerous inter-domain interactions in the AGO-small RNA complex, but the N-coil/Piwi interface is only moderately hydrophobic with several residues calculated to prefer a hydrophilic environment (Figure 1C). For example, Arg28 in human Ago2, equivalent to Arg182 in arabidopsis AGO1, participates in a salt bridge deep inside of the PIWI domain (Figure 1D). This may indicate that the conformation of the N-coil revealed in the crystal structures of AGO-small RNA complexes is but one of several possible conformations of this region. Second, specific residues in the N-coil of Arabidopsis AGO1 (Lys178, Lys185, Gly186 and Lys190) are required for yeast two-hybrid interaction of the N-domain of AGO1 with factors implicated in regulated proteolysis, including the autophagy cargo receptor ATI1^23^. Combining the indication of alternative N-coil conformations, the known instability of unloaded AGO proteins, and the N-coil-dependent two-hybrid interactions with regulated proteolysis factors, we hypothesize that the N-coil is accessible in the unloaded conformation of AGO1, but becomes attached to the Piwi domain as part of the loading process. We further hypothesize that the accessibility of the N-coil in the unloaded form is used for its specific recognition by factors implicated in regulated proteolysis, thereby providing a molecular explanation for the rapid turnover of unloaded AGO1 *in vivo* (Figure 1E). The remainder of this paper describes the results of tests of predictions that follow immediately from this hypothesis for N-coil function.

### The AGO1 Y691E protein mutated in the 5’-phosphate binding pocket is unloaded

Since all tests of our model require manipulation of unloaded AGO1, we first engineered the Y691E mutant of AGO1, mutated in a key residue in the MID domain that coordinates the 5’-phosphate of the 5’-nucleotide of the small RNA and stacks with its nucleobase^11,12^, typically a uracil. In other AGO proteins, the equivalent mutation leads to complete loss of small RNA binding ^20,24^. We used stable transgenic lines in arabidopsis to establish that the Y691E mutant is indeed unloaded. Immunoprecipitation of wild type and Y691E mutant protein followed by polynucleotide kinase-mediated labeling of co-purified small RNA showed that in contrast to AGO1^WT^, no small RNA was bound by the AGO1^Y691E^ protein (Figure 2A). Thus, the AGO1^Y691E^ mutant protein is unloaded. Consistent with this molecular result, expression of the AGO1^Y691E^ mutant protein in the *ago1-3* knockout background failed to rescue the knockout mutant phenotype, demonstrating total loss of function (Figure 2B). Identical results were obtained with the AGO1^Y691E^ protein containing additional mutations in the N-coil and N-domain residues required for ATI1 interaction in yeast (Figure 2A,B).

**Figure 2.**
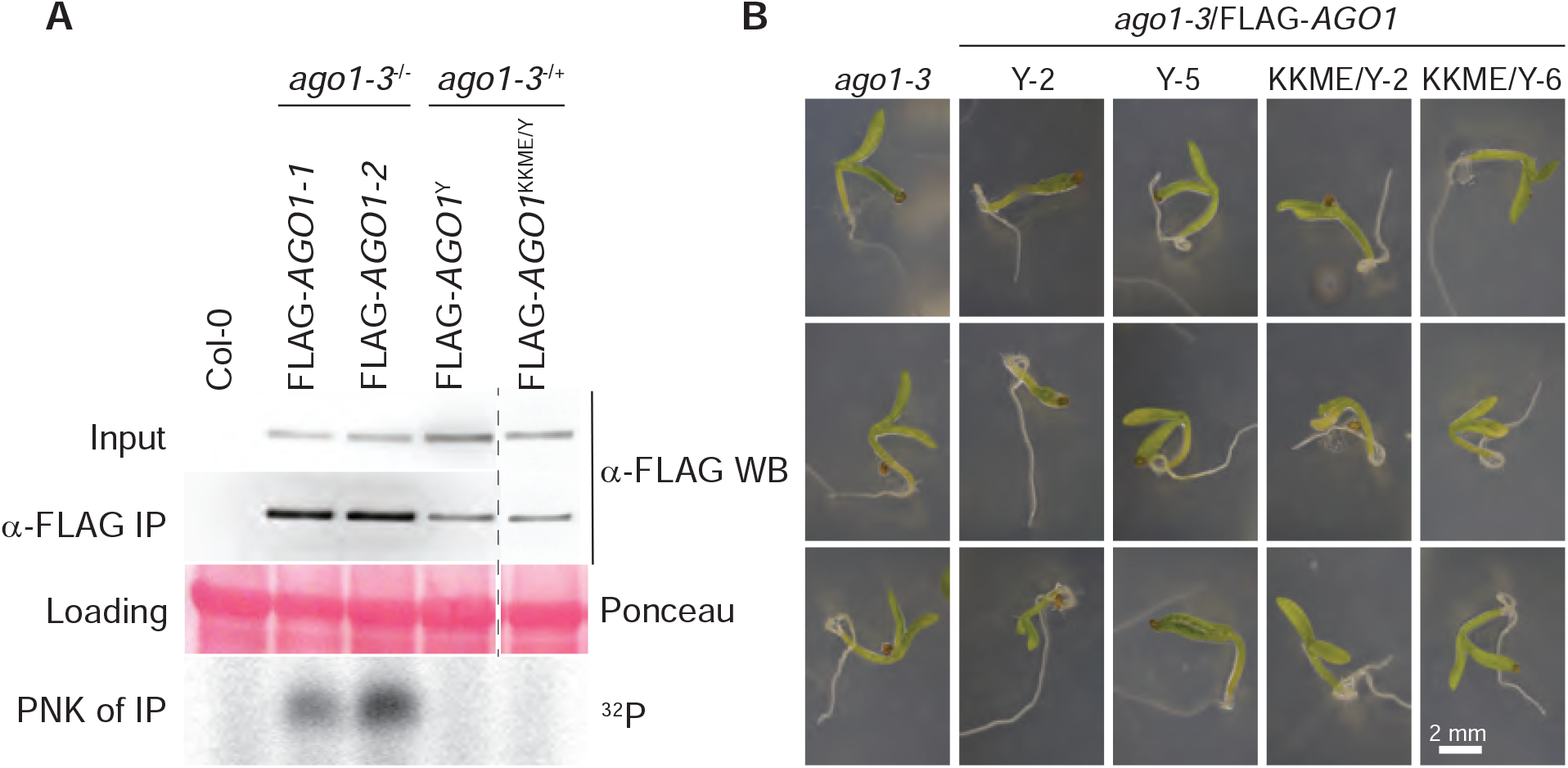
AGO1^Y691E^ does not associate with small RNA. (A) Immunoblot analysis with α-FLAG antibody of total protein and α-FLAG immunoprecipitated protein from inflorescence tissue from the indicated genotypes. The lower panel shows polynucleotide kinase labeling of RNA extracted from α-FLAG IPs. Ponceau stain shows total protein. Y, Y691E; KKME, K185E/K190E/M304E/E307A. Numbers (Y-2, Y-5, KKME/Y-2, KKME/Y-6) denote independent transgenic lines. (B) Images 8 days after germination of homozygous *ago1-3* (null) plants and transgenic plant lines expressing FLAG-AGO1^Y691E^ or FLAG-AGO1^KKME/Y691E^ in the *ago1-3* homozygous background.

### The N-coil is uniquely accessible in unloaded AGO1

We decided to test accessibility of the N-coil in small RNA-bound and free AGO1 using antibody probing. Hence, we raised antibodies against a 16-aa N-coil peptide (F177-C192). This polyclonal antibody specifically recognized the N-coil as demonstrated by its reactivity towards FLAG immuno-purified, denatured AGO1 with a wild type N-coil, but substantial loss of reactivity with FLAG-AGO1 carrying two mutations (K185E/K190E) in the N-coil (Figure 3A-B). We next used this antibody to probe the accessibility of the N-coil in AGO1 in two different ways. First, we observed that binding of equal amounts of N-coil antibody required 10-fold higher amounts of immuno-purified, native, immobilized FLAG-AGO1^WT^ (largely small RNA-bound, see below) than of similarly purified unloaded AGO1 (FLAG-AGO1^Y691E^) (Figure 3B-C). We note that this difference cannot be explained by potentially different patterns of post-translational modifications in the N-coil, because the antibody recognized denatured forms of the two AGO1 variants equally efficiently (Figure 3B). Second, direct immunoprecipitation from total lysates using the N-coil antibody resulted in markedly more efficient recovery (∼10 fold) of FLAG-AGO1^Y691E^ than of FLAG-AGO1^WT^ (Figure 3D), in contrast to the result obtained with FLAG antibodies (Figure 3B). These observations provide clear evidence that the N-coil of AGO1 is preferentially accessible in the unloaded state.

**Figure 3.**
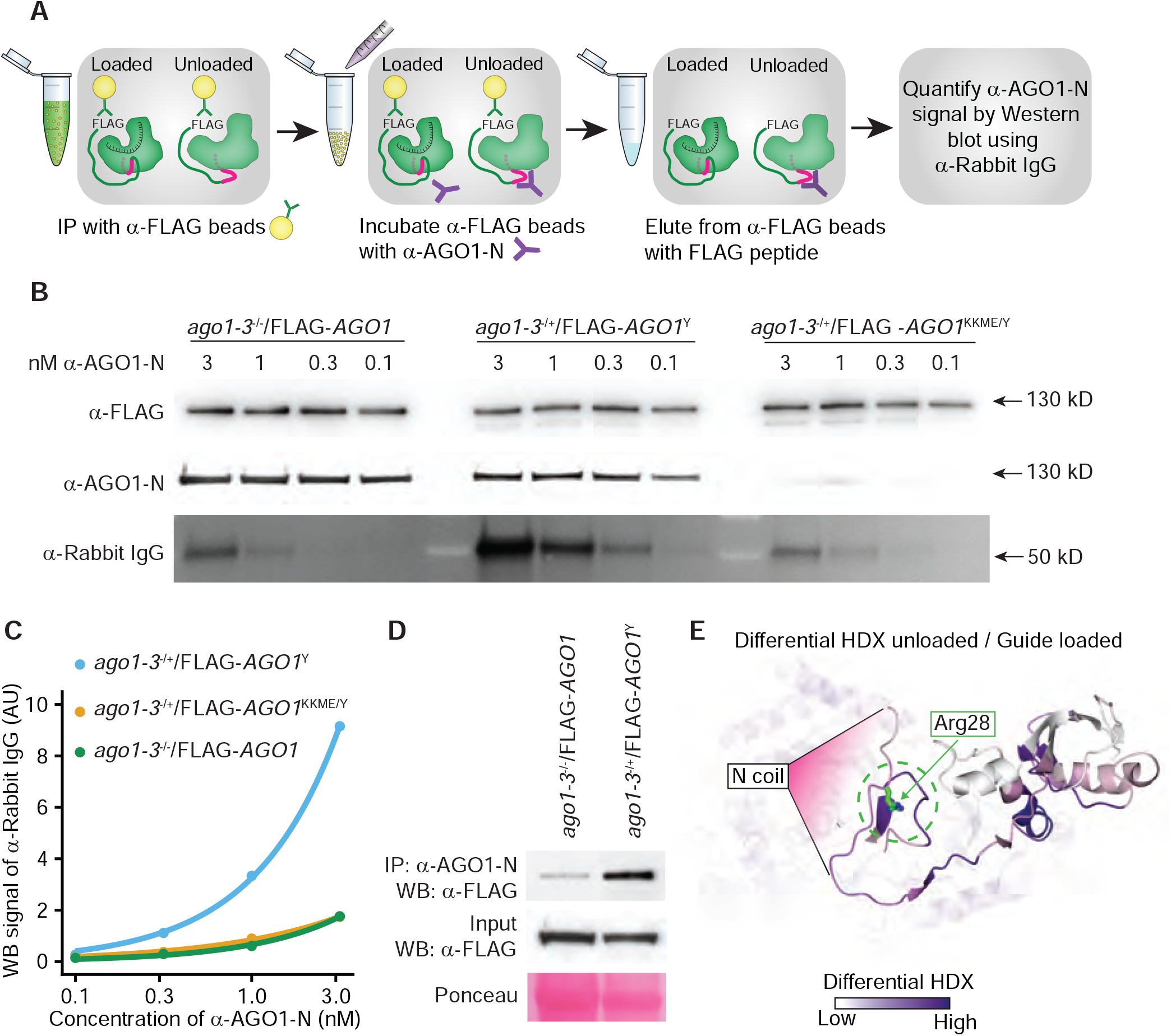
Preferential accessibility of the N-coil in the unloaded state of AGO proteins. (A) Schematic representation of the *in vitro* binding experiment shown in (B) and (C). (B) Protein blots of identical amounts of immobilized, FLAG immuno-affinity purified FLAG-AGO1 variants (WT, Y691E, KKME/Y691E) incubated with the indicated concentrations of AGO1 N-coil antibody (α-AGO1-N) and competitively eluted after extensive washing. Top panel, detection with FLAG antibodies (α-FLAG) to document equal amounts of FLAG-AGO1 variants in each of the binding assays. Middle panel, detection with α-AGO1-N to document specificity of the antibody (compare AGO1^Y691E^ (intact N-coil) to AGO1^Y691E/^KKME (two point mutations (K185E/K190E) in N-coil) and equal reactivity towards loaded and unloaded AGO1 in the denatured state. Bottom panel, detection with anti-rabbit IgG to quantify the amount of α-AGO1-N retained by the immobilized FLAG-AGO1 variants. KKME, K185E/K190E/M304E/E307A. (C) Quantification of the amount of α-rabbit IgG signal in (B) at different α-AGO1-N concentrations. The lines show an exponential fit to the data as a trendline. (D) Immunoprecipitation of FLAG-AGO1^WT^ and FLAG-AGO^Y691E^ with α-AGO1-N antibody from total lysates of plants expressing comparable levels of FLAG-AGO1^WT^ and FLAG-AGO^Y691E^. The Ponceau stain of the lanes containing aliquots of total lysates shows a crop around the large subunit of RuBisCO. (E) Projection onto the *Hs*Ago2 structure of the published fractional differential hydrogen-deuterium exchange (HDX) between unloaded and guide loaded hAgo2^24^. Increasing differences in exchange rate are indicated with progressively darker purple shading. All regions with differential rates of exchange between unloaded and guide-loaded *Hs*Ago2 show higher exchange rate in the unloaded state, most notably areas buried by the N-coil and engaging in interactions with N-coil residue Arg28 in the guide-loaded *Hs*Ago2.

### The N-coil-Piwi interface of human Ago2 exhibits reduced solvent exposure upon small RNA binding

We next wished to assess whether differential N-coil accessibility applies more generally to eukaryotic AGO proteins. To this end, we turned to recently published hydrogen-deuterium exchange data on RNA-free and small RNA-associated human Ago2^25^. Amide protons exchange with the solvent, and their rate of hydrogen-deuterium exchange upon dissolution in D_2_O is, therefore, a sensitive measure of solvent exposure and engagement in hydrogen bonds^26^. Indeed, comparison of RNA-free, guide RNA-associated and guide/target RNA-associated human Ago2 identified regions with differential hydrogen-deuterium exchange rates between the three states^25^. Interestingly, reanalysis of this data with an eye towards the N-coil and interacting residues in the Piwi domain in the loaded conformation showed that regions of slower amide proton exchange in loaded versus unloaded human Ago2 include the region of interest: the N-coil and, in particular, the nearby hydrophilic environment in the Piwi domain caging the guanidium group of the conserved Arg28 side chain in the RNA-bound state (Figure 3E, Figure 1A). Taken together, the antibody probing of arabidopsis AGO1 and the differential hydrogen-deuterium exchange rates in human Ago2 upon loading show that structural flexibility and preferential accessibility of the N-coil in the unloaded state is a recurrent feature of eukaryotic AGO proteins.

### The interaction between ATI1 and the N-domain is direct and involves the N-coil

To test the second prediction that regulated proteolysis factors directly bind the N-coil, we chose the interaction between a fragment of AGO1 comprising the N-coil and the globular N-domain (NcGN) and the N-terminal IDR of the autophagy cargo receptor ATI1^27^ as model system. Using yeast two-hybrid assays, this interaction was shown to depend on the N-coil residues Lys178, Lys185 and Lys190^23^. We first expressed and purified the IDR of ATI1^27^ and wild type and K178E mutant versions of the NcGN of AGO1 fused to His_6_-SUMO (His_6_-SUMO-NcGN^AGO1^, Figure 4A), and used these purified proteins for quantitative binding assays by microscale thermophoresis. This assay uses the distinct thermophoretic mobility of different molecular entities to measure the percentage of complex formed between a fluorescently labeled component, and a partner protein added in increasing concentrations from none (0% complex) to an estimated 100-fold molar excess (∼100% complex)^28^. In our experimental set-up, His_6_-SUMO-NcGN^AGO1^ (WT or K178E) was fluorescently labeled on the unique Cys192 residue close to the N-coil, and the unlabeled IDR of ATI1 was added in increasing quantities. Using this set-up, we showed that the interaction between NcGN^AGO1^ and the IDR of ATI1 is direct, and measured the dissociation constant to be 11±2 µM (Figure 4B). Remarkably, these affinity measurements did not necessitate application of a thermal field to measure differential molecular movement, because the fluorescence intensity itself was sensitive to addition of the IDR of ATI1 (Figure 4C). This observation suggests that in the NcGN^AGO1^-ATI1(IDR) complex, the C192-linked fluorophore is located in a distinct chemical environment compared to the free NcGN^AGO1^, and thus, close to an interaction site. Indeed, when the N-coil mutant K178E was analyzed in the same assay, both the affinity of the ATI1-IDR interaction (47±12 µM), and, especially, the change of fluorescence upon complex formation were substantially reduced (Figure 4B-C). These observations indicate that the N-coil is a direct site of interaction between NcGN^AGO1^ and ATI1-IDR, and that it contributes substantially to the affinity of the interaction.

**Figure 4.**
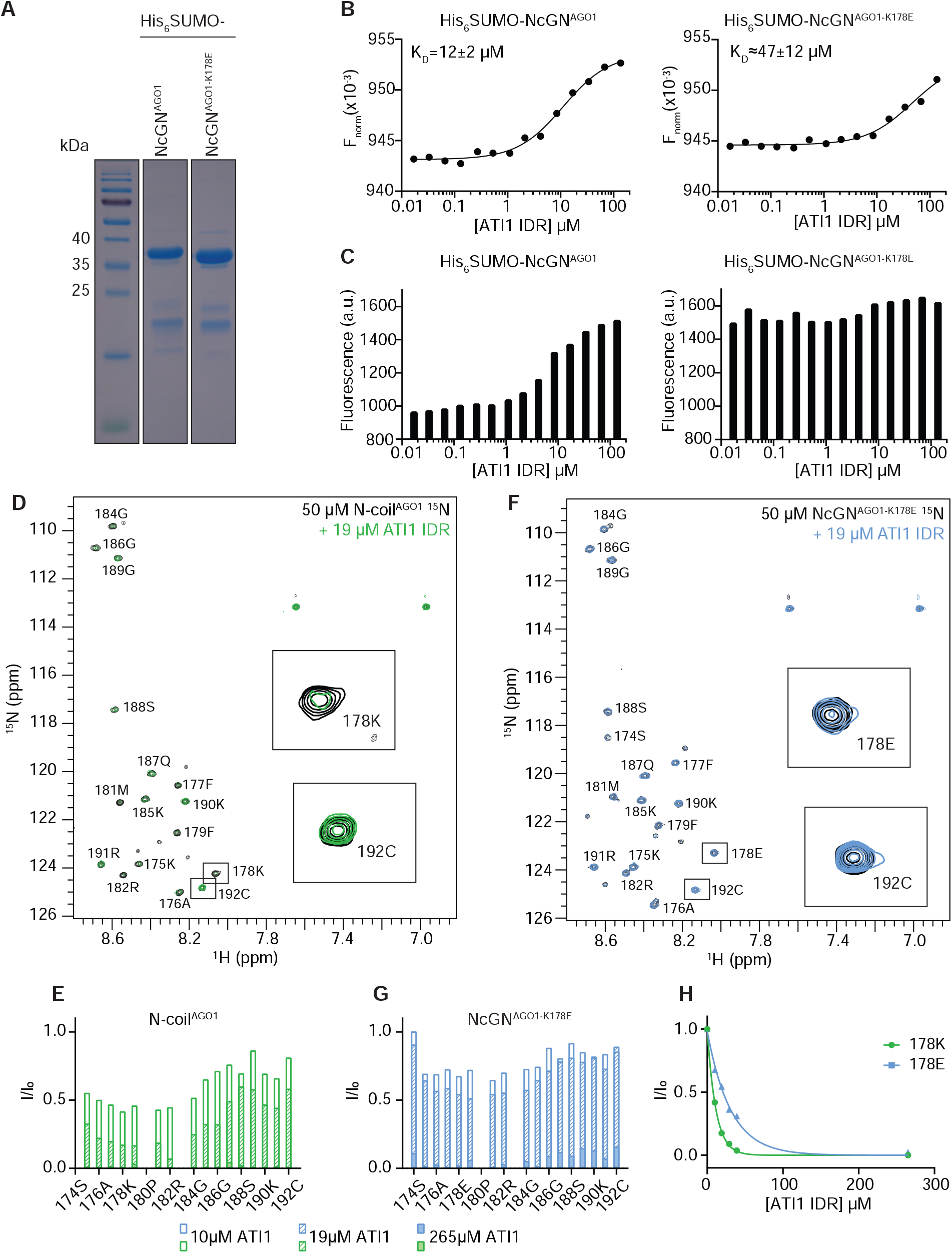
The N-coil is a direct binding site for ATI1 *in vitro*. (A) SDS-PAGE of heterologously expressed His_6_-SUMO-NcGN^AGO1^ protein used for fluorescence labeling and microscale thermophoresis (MST) assays. (B) MST measurements of fluorescence intensity of His_6_-SUMO-NcGN^AGO1^ WT (left) and K178E (right) after application of a temperature gradient. Fluorescence data obtained from the thermophoretic acquisition period were normalized to the initial fluorescence before being fitted to a one-site binding curve with increasing concentrations of ATI1-IDR. The reported binding constants (K_D_) are calculated with standard errors. (C) Simple fluorescence intensity of Cys192-labeled His_6_-SUMO-NcGN^AGO1^ WT (left) and K178E (right) as a function of the concentration of ATI1-IDR. (D, F) ^1^H,^15^N-HSQC nuclear magnetic resonance spectra of 50 μM of ^15^N-labeled AGO1 N-coil WT (left, black) or K178E peptide (right, black), with 19 μM ATI1-IDR added (green with N-coil WT, blue with N-coil K178E). Zooms of contour levels of residues K178 and C192 are highlighted in the boxes. Minor peaks were assigned to a C-terminally truncated version of the AGO1 N-coil peptide whose presence in the sample was verified by MALDI-TOF mass spectrometry. (E, G) Peak intensity plots of ^15^N-labeled AGO1 N-coil peptide (50 μM) upon titration of ATI1-IDR. Peak intensities are normalized to peak intensities observed with free AGO1 N-coil peptide. (H) Peak intensity of residues K178 (for the AGO1 wild type N-coil, green) and E178 (for the K178E N-coil mutant, blue) as a function of the ATI1-IDR concentration. The data points are fitted to an exponential function as a trendline.

### Residues in the N-coil interact directly with ATI1

To show conclusively that the N-coil is a direct interaction site of ATI1-IDR, we expressed and purified an ^15^N-labeled N-coil peptide (172-SSSKAFKFPMRPGKGQSGKRC-192) to perform 2D ^1^H-^15^N heteronuclear single quantum coherence nuclear magnetic resonance (NMR) spectroscopy in the presence and absence of the ATI1-IDR. These experiments showed that the signal intensities from backbone amide N-H nuclei in the N-terminal part of the N-coil (Phe177-Lys185), and, to a lesser extent, the C-terminal Lys190, were strongly reduced upon addition of ATI1-IDR (Figure 4D,E), indicating that these residues participate directly in the interaction. Consistent with the importance of Lys178 for the ATI1-IDR interaction, such loss of signal was markedly less pronounced upon addition of ATI1-IDR to an ^15^N-labeled N-coil peptide containing the K178E mutation, in particular for the amide group of residue 178 (K/E) itself (Figure 4F-H). We conclude that the N-coil is a direct interaction site of the ATI1-IDR, and that the interaction involves contacts along the entire N-coil, including the Lys178, Lys185, and Lys190 residues.

### ATI1 interacts with unloaded AGO1 in vivo and the binding implicates N-coil residues

Having established that the N-coil interacts directly with a protein implicated in regulated proteolysis, we next wished to address if this interaction could occur in the context of full-length AGO1, in particular in the unloaded state. Because heterologous expression of AGO1 failed, even with the use of a variety of expression hosts (*E. coli*, baculovirus/*Sf-9, Schizosaccharomyces pombe*) and affinity/solubilization tags, we used bimolecular fluorescence complementation (BiFC) assays upon transient expression in *Nicotiana benthamiana* for assessment of ATI1 interaction with AGO1^WT^ and AGO1^Y691E^. The close AGO1 paralogue, AGO10, whose NcGN part interacts much less strongly with ATI1 in the two-hybrid assay than AGO1^23^, was used as a control expected to yield substantially less specific signal than AGO1. With AGO1^WT^ fused to the C-terminal part of YFP and ATI1 fused to its N-terminal part (C-YFP-AGO1^WT^ and N-YFP-ATI1), weak, but specific signal was detected in association with the endoplasmic reticulum (ER; Figure 5A), consistent with previous reports of ATI1-localization to ER-associated ATI-bodies, and with peripheral ER-association of AGO1^29,30^. We detected a consistent and statistically significant reduction in intensity of the BiFC signal when the C-YFP-AGO1^K185E/K190E^ mutant containing substitutions in N-coil residues implicated in ATI1 binding was used (Figure 5A-B; Figure S3). We next compared patterns of BiFC signal obtained with C-YFP-AGO1^Y691E^/N-YFP-ATI1 and C-YFP-AGO1^Y691E/K185E/K190E^/N-YFP-ATI1. With both forms, specific signal was detected, and, similar to what we observed with the wild type AGO1, the K185E/K190E N-coil mutations resulted in significantly reduced BiFC signal intensity (Figure 5A-B). These results demonstrate that ATI1 interacts with the unloaded form of AGO1, and that N-coil residues contribute to this interaction *in vivo*. We note that complete disruption of ATI1-AGO1 interaction is not expected upon mutation of N-coil residues, because a previous study showed yeast two-hybrid interactions between ATI1 and AGO1 involving both N-PAZ and MID-Piwi lobes of the protein^31^. We also note that the results show that ATI1 is capable of interacting with unloaded AGO1, and may possibly only interact with the unloaded form, because wild type AGO1 exists as a mixed population *in vivo*, mostly of the RNA-bound form, but also with a minor fraction in the unloaded form (see below). Clearly, however, such possible exclusive interaction with unloaded AGO1 is not established by the BiFC assays.

**Figure 5.**
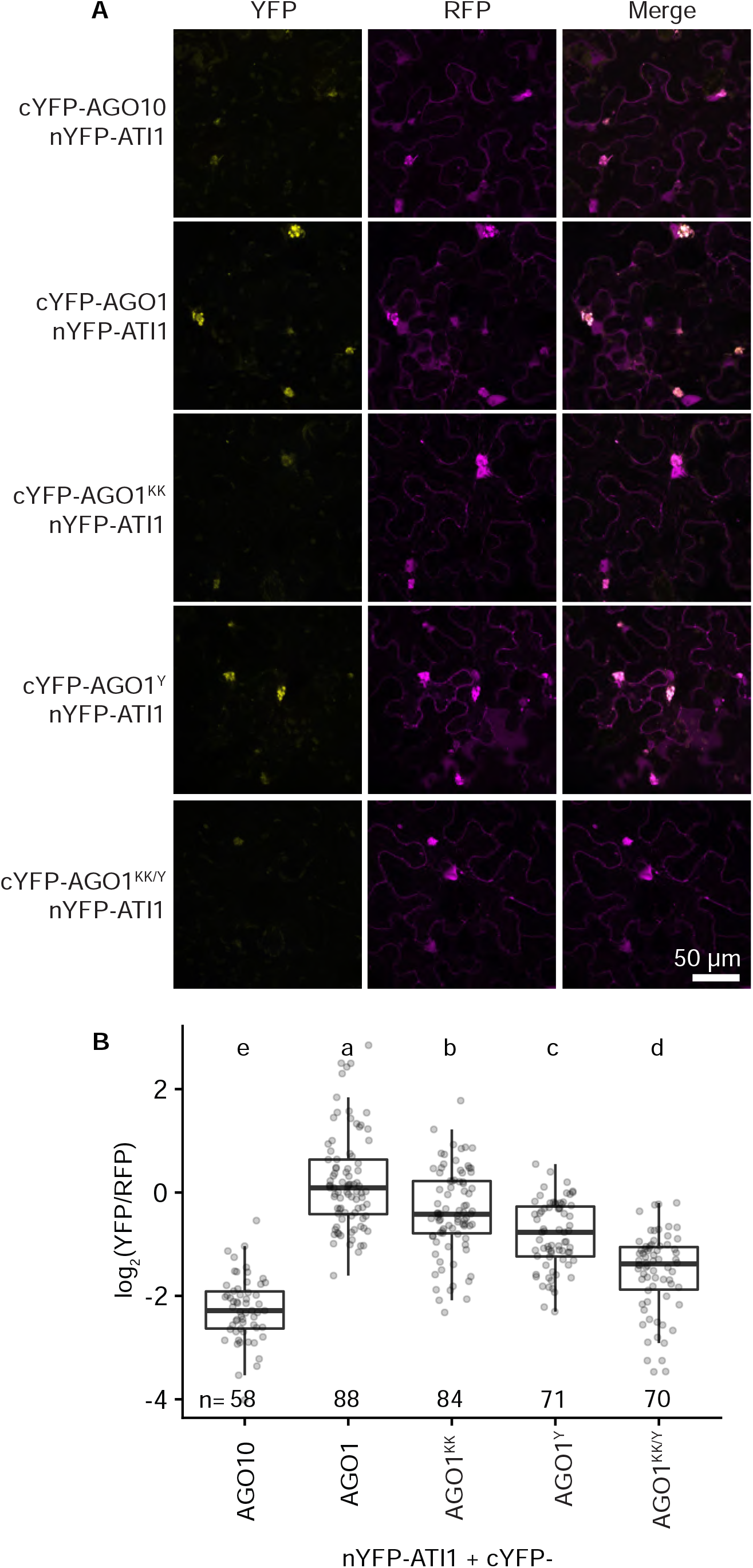
Unloaded AGO1 interacts with ATI1 *in vivo*. (A) Confocal images showing results of bimolecular fluorescence complementation (BiFC) assays to probe the AGO1-ATI1 interaction *in vivo*. The C-terminal half of YFP was fused to AGO10, or to wild type or unloaded (Y691E) AGO1 with either an intact or a mutated (K185E/K190E) N-coil. The N-terminal half of YFP was fused to ATI1. Both fused halves of YFP were expressed from the same plasmid also expressing free RFP. For each BiFC pair, possible interaction is detected by fluorescence from reconstituted YFP. RFP fluorescence serves as a control for transformation. Merged channel images demonstrate the presence of yellow fluorescence only in transformed cells. Scale bar, 50 μm. (B) Quantification of YFP signal normalized to RFP signal in the given number of YFP-positive cells for the interactions shown in (A). Letters indicate groups with statistically significantly different log_2_(YFP/RFP) ratios (p < 0.01; one-way ANOVA with post-hoc Tukey Honestly Significant Difference test). See Figure S3 for image data underlying the quantification in (B). KK, K185E/K190E.

### Defective ATI1 interaction sites in NcGN^AGO1^ reduce the degradation rate of unloaded AGO1

We finally tested the relevance of residues in the NcGN^AGO1^ necessary for ATI1 interaction for degradation of the full-length, unloaded AGO1 protein using pulse-labeling analysis. To maximize chances of measuring an effect *in vivo*, we used the K185E/K190E/M304E/E307A (henceforth, KKME) mutant with residues important for ATI1 interaction as identified by two-hybrid assays mutated both in the N-coil and in the globular N-domain^23^. Thus, stable transgenic lines expressing equal steady-state levels of FLAG-AGO1^WT^, FLAG-AGO1^Y691E^, and FLAG-AGO1^Y691E/KKME^ were selected as material for kinetic analyses. We first considered the possible kinetic pathways of newly synthesized AGO1^WT^ and AGO1^Y691E^. Because AGO1^WT^ can be degraded either prior to or after small RNA loading (Figure 6A), the labeling data must be fitted to a sum of two exponentials, with half-lives corresponding to unloaded (minor fraction, 8.5±1.8%, see Methods) and loaded states (major fraction, 91.5±1.8%, see Methods), respectively. In contrast, labeling of the unloaded AGO1^Y691E^ is expected to follow mono-exponential kinetics. Initial analyses indicated that ^35^S-Met/Cys labeling of entire seedlings resulted in a fairly uniform distribution of label within 15 minutes (Figure S4), but that a classical pulse-chase set-up was not feasible, because incorporation continued after addition of the chase. Since synthesis and degradation rates are equal at steady state, we decided to do pulse-labeling experiments and measure protein half-lives from incorporation rates of ^35^S-label/total AGO1 protein in FLAG immunoprecipitations following addition of ^35^S-Met/Cys label (see Methods). We first compared FLAG-AGO1^WT^ and FLAG-AGO1^Y691E^ lines expressing comparable levels of FLAG-AGO1 at steady state (Figure 6B-C, Figure S5). This analysis showed that wild type AGO1 indeed exhibits two-phase kinetic behavior. Gratifyingly, the half-life of the rapid phase roughly matches that of unloaded AGO1 measured with the Y691E mutant [*t*_1/2_(WT, rapid) = 1.6±0.6 h, *t*_1/2_(Y691E) = 2.5±0.5 h]. We therefore repeated the pulse-labeling experiments with wild type and NcGN interaction site mutants in the unloaded FLAG-AGO1^Y691E^ version, i.e. FLAG-AGO1^Y691E^ vs FLAG-AGO1^Y691E/KKME^.

**Figure 6.**
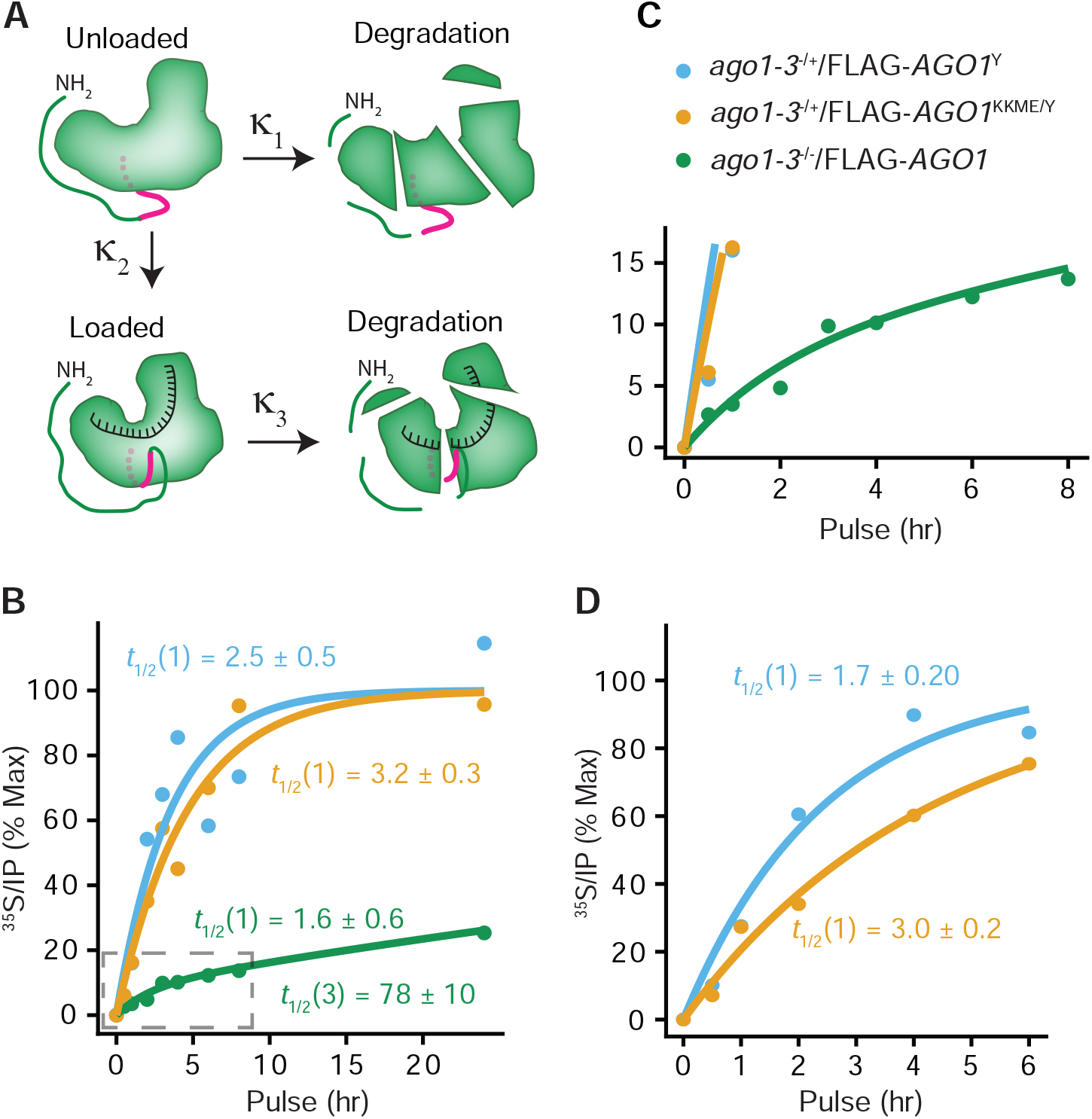
An intact AGO1 N-coil is required for maximal turnover rate of unloaded AGO1 *in vivo*. (A) Kinetic model of the two different fates of newly synthesized AGO1: direct degradation (κ_1_), or small RNA loading (κ_2_) followed by degradation (κ_3_). (B) Quantification of ^35^S-Met/Cys incorporation into FLAG-AGO1^WT^, FLAG-AGO1^Y691E^ (FLAG-AGO1^Y^) or FLAG-AGO1^KKME/Y691E^ (FLAG-AGO1 ^KKME/Y^) as a function of pulse time. The lines represent a fit to an exponential equation for unloadable AGO1 containing the Y691E mutation and a fit to a double exponential equation for the WT AGO1. The half-lives calculated by the fitted models are given in hours +/- standard deviation, as detailed in Methods. KKME, K185E/K190E/M304E/E307A. (C) Zoom in of the dashed area of (B). (D) Result of an independent experiment done as in (C) to verify the slower incorporation kinetics in FLAG-AGO1^KKME/Y691E^ compared to FLAG-AGO1^Y691E^. See Figure S5 for raw data underlying the graphs in Figure 6B-D.

These experiments showed a modest, but reproducible reduction in the degradation rate of the unloaded AGO1 protein with mutations in NcGN residues important for ATI1 interaction (Figure 6B,D; Figure S5; results of two independent experiments shown to convey their quantitative variability). We note that the reduced degradation rate of FLAG-AGO1^Y691E/KKME^ compared to FLAG-AGO1^Y691E^ was not due partial loading of the FLAG-AGO1^Y691E/KKME^ protein, because small RNAs were undetectable in immunopurified fractions of FLAG-AGO1^Y691E/KKME^ and because the mutant showed complete loss of function when expressed in *ago1-3* knockout backgrounds, indistinguishable from FLAG-AGO1^Y691E^ (Figure 2A,B). Hence, we conclude that intact NcGN^AGO1^ interaction sites, including the N-coil, are necessary for maximal turnover rates of unloaded AGO1 *in vivo*.

## DISCUSSION

Our results demonstrate that the N-coil is exposed for interaction preferentially in the unloaded state of AGO1. This is important, because it defines a conformational change that accompanies the AGO loading process, and, thereby, confers distinct protein-protein interaction properties on the small RNA-bound and free states of AGO. In turn, this property offers a molecular explanation of the longstanding observation that unloaded AGO proteins in plants and animals turn over much more rapidly than the small RNA-bound form, as demonstrated here for arabidopsis AGO1. Given the structural conservation of the N-coil in different eukaryotic AGO proteins, in particular its structural flexibility dependent on loading status, it is likely that this same molecular principle operates more generally to confer rapid turnover specifically on unloaded AGO proteins. Of the many questions related to N-coil function that now call for answers, we select two for further discussion. (1) How widespread a phenomenon across AGO proteins is the existence of a structurally flexible, MID-Piwi associated N-coil? In particular, is there evidence for its existence in eukaryotic Piwi-clade proteins and in prokaryotic AGO proteins? (2) Could the N-coil have functional significance in addition to serving as a molecular handle for specific recognition of AGO proteins in the free state?

### An N-coil in Piwi-clade proteins, but not in prokaryotic Ago proteins

Piwi-clade proteins are also likely to exhibit instability in the unloaded state, because their steady-state accumulation is reduced in the absence of cognate piRNAs^32^. Thus, it is a question of interest whether Piwi-clade proteins have an N-coil and what properties it may have. Structures of three Piwi-clade proteins in the piRNA-bound state (silkworm (*Bombyx mori*) Siwi, *Drosophila melanogaster* (Dm) Piwi, and the sea sponge *Ephydatia fluviatilis* (Ef) Piwi) have been reported^33-35^. The Siwi structure shows an N-coil (Ile130 to Gly148, ISILRTRPEAVTSKKGTSG) associated with the MID-Piwi lobe much like in Ago-clade structures (Figure 7A). Thus, N-coil interaction may also be used for molecular recognition of unloaded Piwi proteins, an exciting possibility given that recognition of the unloaded state has significance beyond regulated proteolysis for Piwi proteins. For instance, in insect germline cells, Piwi proteins engage in a so-called ping-pong amplification cycle of piRNAs derived from transposable elements. Ping-pong amplification entails cleavage of a sense piRNA precursor transcript by a mature piRISC containing the Piwi protein Aubergine (Aub) bound to a primary, anti-sense piRNA, and subsequent loading of the sense cleavage fragment onto another Piwi-clade protein, Ago3^36-38^. Ago3 is precluded from primary piRNA association by the protein Krimper that associates with Ago3 specifically in the free state^39^. The molecular determinants of the recognition of the unloaded state of Ago3 by Krimper may include unmethylated Arg residues in the N-terminal IDR of Ago3^39^, but it is now a plausible possibility that the N-coil of Ago3 is an additional such determinant. In contrast, structures of prokaryotic AGO proteins in the nucleic acid-bound state do not show indication of an N-coil^40-43^. Indeed, in prokaryotic AGO proteins, the area in the Piwi domain corresponding to the one surrounding the conserved N-coil residue Arg28 in human Ago2 is markedly different from eukaryotic AGO proteins. In eukaryotic AGO proteins, this region appears optimized for existence in alternative states depending on N-coil/Piwi interaction, because a loop containing the conserved His742 and Asp747 in *Hs*Ago2 is held in place by the buried Arg28 N-coil residue. We note that this conformation also involves a characteristic cluster of His residues (Figure 7B) whose degree of protonation may favor one state over the other. In contrast, in prokaryotic AGO proteins, this region is structurally fixed (Figure 7C), consistent with the absence of a flexible, Piwi-interacting N-coil.

**Figure 7.**
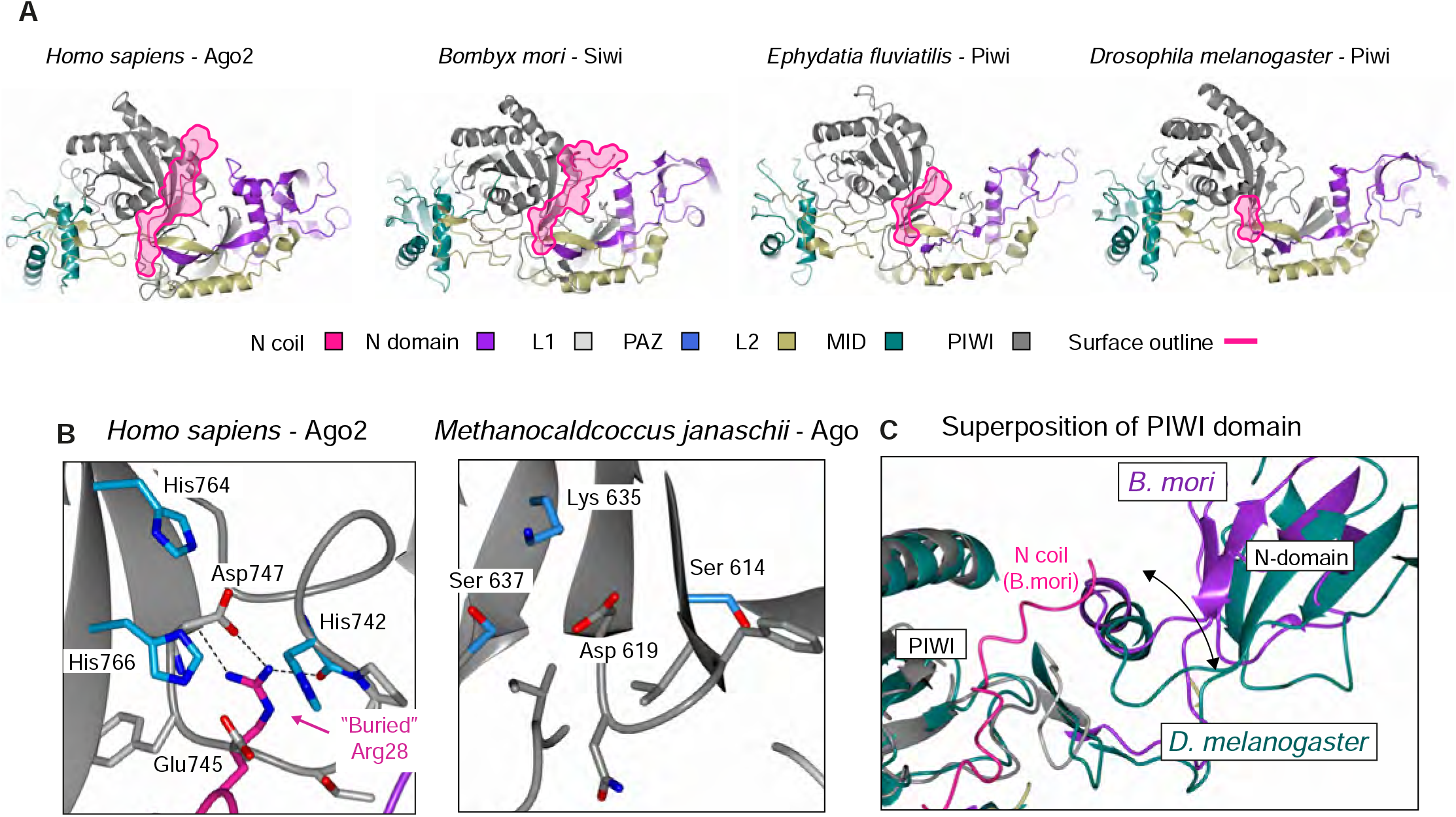
Piwi-clade, but not prokaryotic, AGO proteins have a structurally flexible N-coil. (A) Structural overview of *Hs*Ago2 and Piwi proteins, outlining their N-coils in pink. The *Ephydatia fluviatilis* and *Drosophila melanogaster* Piwi proteins harbour N-coil truncations, probably generated by proteolysis during sample preparation for structural studies^33,34^. The structural representations are based on the published coordinates (*Hs*Ago2, PDB: 4F3T^10^; *Bm*Siwi PDB: 5GUH^32^; *Ef*Piwi, PDB: 7KX7^34^; *Dm*Piwi PDB: 6KR6^33^). (B) Close-up of region in the Piwi domain in eukaryotic *Hs*Ago2 (PDB: 4F3T^10^) that cages the buried N-coil Arg28 side chain in the small RNA-bound conformation. Contacts between Arg28 and the loop containing the conserved His742, Glu745 and Asp747 residues are shown by dashed lines. A cluster of histidines formed by residues in β-strands of the Piwi-domain (His764, His766) and in the Arg28-interacting loop (His742) is highlighted in blue. (C) Close-up of the region in the Piwi domain of the prokaryotic *Mj*Ago (PDB: 5G5T^42^) that corresponds to the region of *Hs*Ago2 shown in (B). (D) Secondary structure matching of the *B. mori* and *D. melanogaster* Piwi domains reveals a more relaxed state of the *D. melanogaster* Piwi molecule (green) in which the structurally fixed N-coil seen in *B. mori* Siwi is absent.

### Relevance of N-coil-Piwi interactions for the mature RISC

While accessibility of the N-coil for protein-protein interaction in the unloaded state can be exploited for specific recognition of the unloaded form, it is also relevant to consider possible functions of the N-coil in the mature RISC. Since the N-coil-Piwi interactions seen in structures of small RNA-bound AGO proteins occur exclusively in the loaded conformation as judged by the antibody probing results of arabidopsis AGO1 and differential hydrogen-deuterium exchange rates in human Ago2, it is possible that N-coil-Piwi interactions are important to reach the compact conformation of mature RISC. Two observations are compatible with this proposition. First, the N-coil is clearly important for RISC function because two point mutations in this region have been recovered in forward screens for defective miRNA function in arabidopsis: *ago1-38* (G186R) that exhibits relatively weak loss of function^44^, and *ago1-55* (G189E) that exhibits stronger loss of AGO1 function^45^. The exact molecular defects of these point mutants have not been defined, however, and it remains possible that RISC activity rather than loading could be affected in these mutants. Second, closer inspection of the available Piwi-clade structures reveals intriguing differences. While the Siwi protein analyzed has a full N-coil as discussed above, *Ef* Piwi and *Dm* Piwi harbor N-terminal truncations, presumably due to proteolytic cleavage prior to structural characterization^34,35^, such that the analyzed *Ef* Piwi protein has a partial N-coil, and *Dm* Piwi no N-coil at all (Figure 7A). Strikingly, the compactness of the Piwi:piRNA complexes increases progressively from the *Dm* Piwi structure (no N-coil present) to the Siwi structure (full N-coil present) (Figure 7A,D), lending support to the idea that the N-coil/MID-Piwi lobe association may be an important step in reaching the final, compact conformation of mature RISC across both Piwi and Ago clades. Finally, we note that a key functional importance of the N-coil and the *β*-strands connecting it to the rest of the N domain has been observed in human Ago proteins where substitution of this part of the catalytically inactive human Ago3 by the corresponding part of the catalytically active Ago2 is sufficient to confer activity on Ago3^46^. This observation suggests that the relevance of the N-coil interactions with the Piwi domain may not be limited to the loading process, but may also be important for the extensive conformational changes required to reach the catalytically active conformation of AGO proteins^47^.

## MATERIALS AND METHODS

### Plant material and growth conditions

The Columbia-0 (Col-0) *Arabidopsis thaliana* accessions were used. *ago1-3* was described previously^48^. Seeds were sterilized as described in [^49^] prior to being sown on Murashige-Skoog (MS) agar medium (4.3 g/L MS salts, 0.8 % agar, 1% sucrose) where they were grown for 10 days at constant temperature (21 °C) and a 16h light (80 µE m^-2^ s^-1^)/8h darkness cycle.

### Antibodies

Anti-FLAG® M2 Affinity Gel and monoclonal anti-FLAG® M2-Peroxidase (HRP) were purchased from Sigma, anti-HSP70 from Agrisera, and anti-Rabbit IgG (HRP) from Sigma. The anti-AGO1-N-coil antibody was affinity purified by Eurogentec from sera collected from a rabbit immunized with a 16-amino acid peptide from the N-coil of AGO1 (F177-C192, H-FKFPMRPGKGQSGKRC-NH_2_), synthesized by Schafer-N, Copenhagen, Denmark. The cysteine corresponding to C192 in AGO1 was used for KLH coupling prior to immunization.

### Site-directed mutagenesis

Site-directed mutagenesis was performed according to the Quickchange protocol (Stratagene) using Phusion High-Fidelity DNA Polymerase (NEB) and primers containing the appropriate mutations (Supplemental Table 2). The reaction mix was subsequently digested with DpnI (Thermo Scientific) to remove methylated template DNA before transformation to DH5*α* competent *E. coli* (NEB).

### Plant transformation and selection

Transgenic lines were generated by floral dip transformation with *Agrobacterium tumefaciens* GV3101^50^. The FLAG-AGO1^Y691E^ and FLAG-AGO1^KKME/Y691E^ (K185E/K190E/M304E/E307A/Y691E) mutations were introduced to pCAMBIA3300U AGO1P:FLAG-AGO1-AGO1T previously described in [^51^] using site-directed mutagenesis (primers listed in Supplemental Table 2). FLAG-AGO1^Y691E^ and FLAG-AGO1^KKME/Y691E^ were transformed into *ago1-3/+* lines. Primary transformants (T1) were selected on MS agar medium supplemented with 100 µg/ml ampicillin and 7.5 µg/ml glufosinate ammonium (Fluka). Transformed plants were transferred to soil after approximately 10 days of growth on selective medium. For every construct, lines with a single T-DNA locus and comparable protein expression levels were selected in T2, and descendants homozygous for the insertion were identified in T3.

### Genotyping

Genomic DNA from aerial tissue of rosette leaves was genotyped using Phire Plant Direct PCR Kit (Thermofischer). The primers used for genotyping are listed in Supplemental Table 1.

### PNK labeling of AGO1 bound sRNA

RNA was extracted directly from FLAG IP beads by addition of TRI-Reagent (Sigma) as described above except that RNA precipitation was performed in the presence of 20 μg of glycogen. The RNA was 5’-end labeled with [γ-^32^P]-ATP by T4-polynucleotide kinase according to manufacturer (Fermentas). Unbound [γ-^32^P]-ATP was removed by MicroSpin G-25 columns (GE Healthcare). The RNA was separated by electrophoresis in a denaturing 18 % polyacrylamide gel. The gel was exposed to a phosphor screen and imaged on a Typhoon FLA 7000 scanner (GE Healthcare).

### Protein expression and purification

Purification of ATI1N M1-E180 was carried out as previously described ^27^. AGO1 NcGN (S172-Y335) CDS was cloned into a modified pET24a vector encoding His_6_-SUMO N-terminally. The K178E substitution was generated by site-directed mutagenesis. An additional three substitutions (L318A/L321A/L325S) were made by site-directed mutagenesis to enhance solubility (primers listed in Supplemental Table 1). Plasmids of His_6_-SUMO-tagged AGO1 NcGN and His_6_-SUMO-AGO1 NcGN^K178E^ were transformed into *Escherichia coli* BL21 (DE3 7tRNA) Rosetta cells and grown over night at 37 °C in 50 ml LB media supplemented with 50 μg/ml kanamycin and 33 μg/ml chloramphenicol. 2 ml of culture were added to 1 liter LB medium supplemented with 50 μg/ml kanamycin and 33 μg/ml chloramphenicol and grown at 37°C, 150 rpm to an OD_600_ of 0.6-0.8 before induced with 0.5 mM isopropyl β-D-1-thiogalactopyranoside (IPTG) for expression o/n at 18°C, 150 rpm. Cells were harvested by centrifugation at 8,000 x g for 15 min, 4 °C and resuspended in 25 ml cold lysis buffer (25 mM tris-HCl, 10 mM imidazole, 300 mM NaCl pH 8, 1mM TCEP) supplemented with 1 tablet complete protease inhibitor EDTA-free (Roche) before lysed using a French Press (>10.000 psi). The cell lysate was cleared by centrifugation at 30,000 x g for 30 min, 4 °C and the supernatant was passed through a 0.45 μm filter before incubated on 2.5 ml Ni-NTA agarose beads for 1 h at 4°C. The bead suspension was transferred to an empty gravity column and washed with 50 ml 25mM Tris-HCl, 20mM imidazole, 300mM NaCl pH 8, then eluted in 5 ml 25mM Tris-HCl, 300mM imidazole, 150mM NaCl pH 8 and subsequently dialyzed o/n against buffer depleted for imidazole (20 mM Tris-HCl pH 8, 150 mM NaCl, 1 mM β-ME). His_6_-SUMO-AGO1 NcGN was further purified by application on a Superdex 200 16 600 (GE Healthcare) size exclusion chromatographic column connected to an Äkta Purifier system (GE Healthcare) in 20 mM Tris-HCl pH 7.2, 1 mM KCl, 0.05 mM NaCl, 0.01 mM MgCl_2_, 1 mM TCEP. Eluting protein was detected by measurement of absorbance at 280 nm, and fractions of 1 ml were collected and analyzed by SDS-PAGE. Fractions containing His-SUMO-AGO1 were kept at 4°C and used fresh.

For expression of AGO1 N-coil, a cDNA encoding residues S172-C192 was cloned into a pGEX4t1 vector encoding a GST-tag (primers listed in Supplemental Table 1) and transformed into *Escherichia coli* BL21 (DE3 7tRNA) Rosetta cells. For production of the stable isotope-labeled AGO1 N-coil, a culture of the expression strain was added to 1 L of freshly prepared M9 minimal media (22mM KH_2_PO_4_, 42mM Na_2_HPO_4_·2H2O, 17mM NaCl, 1mM MgSO_4_) supplemented with 1:1000 M2 trace element solution, 0.4 % (v/v) glucose, 18 mM NH_4_Cl-^15^N and 100 μg/ml ampicillin and 33 μg/ml chloramphenicol and grown to OD_600_ of 0.6-0.8 before induction with 1mM IPTG for overnight expression at 18°C. Cells were harvested by centrifugation at 8,000 g 15 min at 4 °C and lysed by 2 passages through a French press (>10,000 psi) in ice cold 1x PBS (137 mM NaCl, 2.7 mM KCl, 10 mM Na_2_HPO_4_, and 1.8 mM KH_2_PO_4_, pH 7.4) with 1 mM TCEP and 1 tablet complete EDTA-free protease inhibitor (Roche). The lysate was cleared by centrifugation at 30,000 g for 30 min at 4 °C and subsequently filtered through a 0.45 μm filter before incubation with 2.5 ml glutathione-conjugated sepharose beads (GE Healthcare) for 1h at 18 °C. The beads were packed in an empty gravity column and washed with 50 ml cold 1xPBS. AGO1 N-coil was liberated from the GST-tag by on-bead cleavage o/n at 4 °C with His_6_-TEV in estimated 1:50 molar ratio in 50mM Tris-HCl, 50mM NaCl, 0.5mM EDTA, 1.75mM TCEP, pH 7.2. The flow-through was collected and His_6_-TEV was captured on 0.1 ml Ni-NTA agarose beads for 1h at 4°C. AGO1 N-coil was further purified by reversed-phase chromatography using a 17.35ml Zorbax 300 Å Stable-bond C18 column connected to an Äkta Purifier HPLC system. Trifluoroacetic acid (TFA) was added to the AGO1 sample to a final concentration of 0.1% and centrifuged at 20,000 g for 15 min to remove insoluble aggregates prior to application. The column was equilibrated with 100% buffer A (0.1% TFA [v/v] in MQ water) and AGO1 N-coil was eluted by a gradient into 100% buffer B (70 % [v/v] acetonitrile, 0.1% TFA [v/v] in MQ water) over 10 CV using a flow rate of 3 ml/min. 2 ml fractions were collected by A_220_ detection. AGO1-containing fractions were collected, flash-frozen in nitrogen and subsequently lyophilized overnight. The samples were resuspended in milli-Q water, adjusted to pH 6.6, before again being flash-frozen and lyophilized o/n and stored at -20 °C. Aliquots of AGO1 N-coil-containing fractions were pH-adjusted by the addition of TA30 (30:70 [v/v] ACN:0.1%TFA) and verified by MALDI-TOF mass spectrometry on an α-cyano-4-hydroxycinnamic acid (HCCA)-matrix.

### Microscale thermophoresis (MST)

All MST data was recorded on a Monolith NT.115 (NanoTemper Technologies) at 25 °C with medium MST-power exciting the fluorophore in the red excitation channel. The buffer composition was 20mM tris-HCl pH 7.2, 1mM KCl, 0.05mM NaCl, 0.01mM MgCl_2_, 1mM TCEP, 0.05 % [v/v] tween20. 20 µM His_6_-SUMO-AGO1-NcGN was orthogonally labeled on Cystein-192 using the Monolith NT™ Protein Labeling Kit Red – Maleimide (NanoTemper Technologies). The labeling was performed according to the manufacturer’s instructions. From a stock solution of 271 μM ATI1 IDR, a 1:1 dilution series was prepared and subsequently mixed 1:1 with SUMO-AGO1 NcGN keeping the concentration and hence the fluorescence constant. The protein series was loaded into Monolith NT.115 Premium Capillaries (NanoTemper Technologies) and thermophoresis was measured using MO.Control v1.6.1 software. The raw data recorded at 0.5-1 sec of thermophoresis was retrieved and analyzed with GraphPadPrism 8.0 fitting by the law-of-mass-action

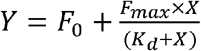

where *Y* is the fluorescence at binding, *F*_*0*_ is the fluorescence when no complex has formed, *F*_*max*_ is the fluorescence at maximum binding, *X* is the concentration of ATI1 IDR, and *K*_*d*_ is the binding constant. All fluorescence intensities were normalized to the initial fluorescence prior to fitting.

### NMR spectroscopy

NMR spectra were recorded on Bruker Avance III 600 MHz or 750 MHz spectrometer equipped with TCI cryoprobes. Spectra were recorded at 5 °C in buffer composition 20mM Tris-HCl pH 6.6, 100mM KCl, 5mM NaCl, 1mM MgCl_2_, 3mM TCEP, 10% (v/v) D_2_O, 250 μM sodium 2,2-dimethyl-2-silapentane-5-sulfonate (DSS). For titration experiments, ^1^H-^15^N-HSQC spectra were recorded of 50 µM ^15^N-labeled AGO1 N-coil in the absence or presence of increasing concentrations of up to 265 µM ATI1 IDR. The intensity ratios of 178K and 178E for different concentrations of ATI1 IDR were fitted as an exponential function in GraphPadPrism 8.0

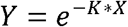

### Assignment of NMR spectra

Assignment spectra were recorded on a Varian Inova 750 MHz magnet equipped with a Bruker Avance III console and a Bruker TCI cryoprobe. Backbone amide resonances of free 1.4 mM AGO1 N-coil and 4.5 mM AGO1 N-coil^K178E^ were assigned manually in CCPNmr Analysis 2.2.4 ^52^ using ^15^N-TOCSY-HSQC and ^15^N-NOESY-HSQC spectra ^53-56^.

### In vitro binding assay of AGO1 variants to the N-coil antibody

1.6 g of inflorescence tissue from the indicated genotypes were ground in liquid nitrogen and extracts were prepared in 6.4 ml IP buffer (50 mM Tris-HCl, pH 7.5, 150 mM NaCl, 10% glycerol, 5 mM MgCl_2_, and 0.1% Nonidet P-40, 4 mM DTT) supplemented with 2x Complete EDTA free protease inhibitor (Roche). Samples were spun at 30,000 *g* for 10 min at 4°C and the cleared lysate was incubated with 40 µl anti-FLAG agarose bead slurry (Sigma) for 30 min at 4°C. The beads were washed three times in IP buffer and divided into four Eppendorf tubes for each genotype. The anti-FLAG beads were incubated with 50 µl of IP buffer containing the indicated concentrations of anti-AGO1N antibody for 20 min at 4°C. The beads were washed three times in IP buffer and eluted twice in 20 µl IP buffer with 100 ng/µl FLAG-peptide (Sigma) for 15 min at 4°C. The samples were heated for 5 min at 75°C in Laemmli buffer, subjected to SDS-PAGE and blotted onto a nitrocellulose membrane. The FLAG-AGO1 proteins were detected with anti-FLAG-HRP (1:4000 [Sigma] in 5 % milk in PBS-T [0.05% tween20]) or anti-AGO1-N-coil (1:1000 in 5% milk in PBS-T) followed by anti-Rabbit antibody (1:10,000 [Sigma] in 5% milk in PBS-T). The anti-AGO1-N-coil antibody was detected with anti-Rabbit antibody conjugated to HRP (1:2,000 [Sigma] in 5% milk in PBS-T). The intensity of the signal from retained anti-AGO1-N-coil antibody was quantified using Image J and fitted using Nonlinear Least Square (NLS) in R (version 4.2.0) with an exponential equation:

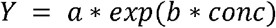

### Immunoprecipitation using the AGO1 N-coil antibody

30 µl pf Protein A-agarose beads (Sigma) were washed three times in PBS and incubated with 500 µl PBS with 7.5 µg of anti-AGO1-Ncoil antibody (1:20 dilution of the fraction affinity purified against the N-coil peptide) for one hour at 4°C and then washed in PBS. In parallel, 200 mg of inflorescence tissue of each genotype were ground in liquid nitrogen and extracts were prepared in 800 µl IP buffer supplemented with 2x Complete EDTA free protease inhibitor (Roche). Samples were spun at 17,000 *g* for 10 min at 4°C and transferred to a new tube. 15 µl of Agarose A beads with anti-AGO1-Ncoil antibody were added to the samples and incubated for 30 min at 4°C. The beads were washed three times in 500 µl IP buffer and heated for 5 min in Laemmli. After SDS-PAGE and blotting onto a nitrocellulose membrane, the FLAG-AGO1 proteins were detected with anti-FLAG-HRP antibody (1:4000 [Sigma] in 5 % milk in PBS-T). The membrane was developed as described above with homemade ECL or SuperSignal™ West Femto Maximum Sensitivity Substrate (Thermo Fisher) and images taken with a Sony A7S camera.

### Bimolecular Fluorescence Complementation (BiFC) assays

BiFC was done using the 2in1 vector system which expressed both the YFPn and YFPc tagged protein as well as free RFP from the same vector^57^. ATI1 was amplified from oligo (dT) primed arabidopsis cDNA and subcloned into pDONR221 P3P2 entry vector with the BP clonase II kit (Invitrogen) and similarly, AGO10 and AGO1 were subcloned into pDONR221 P1P4 (primers listed in Supplemental Table 1). The AGO1^K185E/K190E^ and AGO1^Y691E^ mutations were introduced using site directed mutagenesis (primers listed in Supplemental Table 1). Transfer of the Gateway cassette from these two entry vectors to the pBIFCt-2in1-NN destination vector ^57^ was performed through LR clonase II kit (Agilent).

*Agrobacterium tumefaciens* transformed with the pBIFCt-2in1-NN vector^58^ containing the YFPn- and YFPc-fused cDNAs were infiltrated at an OD_600_ of 0.5 into leaves of *Nicotiana benthamiana* defective in co-suppression achieved by RNAi of *RDR6*^59^. Three days post infiltration, epidermal cells were observed for fluorescence emission using a Zeiss LSM700 confocal laser scanning microscope with a C-Apochromat 63x and a 20x objective. YFP and RFP were excited using laser light of 488 nm 555, respectively. The emitted light was captured with preset filters of the microscope software. All images are shown as z-stack projections of 7-9 slices covering 18-23 µm. Quantification was done on eight z-stack images from individual leaf discs of each construct. The YFP and RFP signal from distinct interaction sites were quantified using Photoshop and the ratio was plotted on a Log2 scale. Log2 transformation was done to obtain normal distribution for the differences in YFP/RFP ratios to be tested statistically using one way ANOVA followed by post hoc Tukey HSD multiple comparison.

### Pulse labeling

Seedlings were grown for 7 days on MS agar under a 16h light/8h darkness light regime. For each time point, 8 seedlings were transferred to 1.5 ml liquid MS in a 12-well culture plate and grown for two more days. EasyTag™ Express35S Protein Labeling-Mix (Perkin Elmer) was then added to a final count of 50 µCi/ml and at given time points, seedlings were briefly washed in water and flash frozen in liquid nitrogen. The tissue was ground to a fine powder before it was solubilized in 500 µl IP buffer (50 mM Tris-HCl pH 7.5, 150mM NaCl, 10 % glycerol [v/v], 5mM MgCl_2_, 0.1% Nonidet P40 [v/v], 4mM DTT, 2 tablets complete EDTA-free protease inhibitor (Roche)) followed by 5 min incubation at 4°C. The lysate was cleared by centrifugation at 16,000 *g* for 5 min at 4 °C before it was immobilized on anti-FLAG (Sigma) agarose bead slurry for 30 min at 4 °C. The beads were transferred to an empty gravity column and washed with 1.5 ml IP buffer. Immunoprecipitated protein was released from the beads in Laemmli buffer by heating at 75 °C for 5 min.

### Pulse quantification

After Western blot development, the nitrocellulose membranes were exposed to a storage phosphor screen and the screen was imaged using a Typhoon FLA 7000 scanner (GE Healthcare). Total lysate and IP Western blot and ^35^S signal were quantified using Image J and ^35^S signal/IP Western blot signal were plotted against pulse time. Due to the unstable nature of the unloaded FLAG-AGO1^Y691E^ *in vitro*, the radioactive signals of the degradation fragments are added to the signal of the intact FLAG-AGO1^Y691E^ protein signal, except for a band identified as HSP70. For all genotypes except FLAG-AGO1^WT^ the pulse data was fitted to an equation of the type,

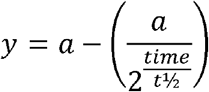

where *t*_1/2_ is the half-life of the immuno-precipitated protein and *a* is the maximum ^35^S value of radioactivity per protein that the sample will converge to when the pulse has reached equilibrium, i.e. when all unlabeled protein has been degraded and the newly synthesized proteins are radio-labelled. Subsequently, the pulse values were adjusted to converge to the value 100 to facilitate comparison between samples and experiments and the data refitted to the equation,

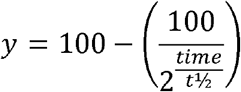

The degradation of FLAG-AGO1^WT^ can be described as a sum of two exponential decays; one for the unloaded fraction and one for the small RNA loaded fraction. Since the steady state protein levels of FLAG-AGO1^WT^ and FLAG-AGO1^Y691E^ and FLAG-AGO1^Y691E/KKME^ are similar, the maximum ^35^S/IP value of FLAG-AGO1^WT^ will converge to the same value as the FLAG-AGO1^Y691E^ and FLAG-AGO1^Y691E/KKME^. Therefore, the FLAG-AGO1^WT^ pulse data can be fitted to an equation of the type,

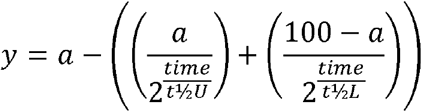

where *a* is the percentage of unloaded AGO1 at steady state, *t*_1/2_*U* is the half-life of the unloaded fraction and *t*_1/2_*L* is the half-life of the loaded fraction. The data analysis was done using Nonlinear Least Square (NLS) in R (version 4.2.0).

### Structural analyses

The alignment of N-coil residues between human Ago2 and plant AGO1 did not require alignment tools. The intra-molecular interactions between the N-coil and the remaining part of *Hs*Ago2 was calculated using the web-service PDBe PISA at the European Bioinformatics Institute (http://www.ebi.ac.uk/pdbe/prot_int/pistart.html)^60^ by treating the N-coil as a separate polypeptide chain. The calculations were done using the human Ago2 structure (PDB 4F3T) and buried surface area (Å) and solvation energy estimates (kcal/mol) are presented for each N-coil residue.

The Fractional Uptake Difference in hydrogen/deuterium exchange between empty and guide RNA-bound human Ago2 was presented on the human Ago2 structure (PDB 4F3T). For this a Pymol script was generated using the available data set ^25^ (MSV000088648 at https://massive.ucsd.edu) at the 5 minute timepoint and the DECA software ver1.12 ^61^. All the presented structural models were made using CCP4mg ^62^.

## ACKNOWLEDGEMENTS

Theo Bølsterli and his team are thanked for plant care. Signe A. Sjørup is thanked for assistance with MALDI-TOF experiments and purification of proteins. We thank Laura Arribas-Hernández and Diego López-Marquez for critical reading of the manuscript. This work was supported by a project grant (NNF17OC0029194) from the Novo Nordisk Foundation, and a Consolidator Grant (ERC-2016-CoG 726417 PATHORISC) from the European Research Council, all to PB. The work was also supported by the Novo Nordisk Foundation Challenge program (REPIN; grant number NNF18OC0033926 to BBK). The NMR spectra were recorded at cOpenNMR, an infrastructure initiative supported by the Novo Nordisk Foundation grant number NNF18OC0032996.

## AUTHOR CONTRIBUTIONS

SB constructed plant lines expressing unloaded FLAG-AGO1 and its combinations with N-coil mutants in *ago1-3* knockout backgrounds, conducted bimolecular fluorescence complementation analyses, pulse-labeling experiments and resulting data analysis, developed the AGO1 N-coil antibody and conducted binding experiments and subsequent data analysis. IMZS expressed and purified recombinant proteins, conducted MST assays with SB and NMR experiments with AP, and participated with SB in preliminary tests and AGO1 binding experiments with a lower-affinity N-coil antibody not used in the final experiments reported here. CP conducted analysis of structures and suggested that the N-coil may adopt conformations different from the one seen in crystal structures of *Hs*Ago2. BBK helped design NMR experiments, and, together with AP, supervised all work and data analysis related to those experiments. PB designed and supervised the study and wrote the manuscript with contributions from all authors.

## DATA AND CODE AVAILABILITY

NMR data have been deposited at the Biological Magnetic Resonance Bank (BMRB) under BMRB entry 51660. The study did not develop new code. Standard procedures for mathematical and statistical analysis of data are described in Methods.

## SUPPLEMENTAL MATERIAL TITLES AND LEGENDS

**Figure S1. Alignment of *Dm*Ago1 with *At*AGO1 and *Hs*Ago2**.

The alignment highlights the position of the Lys514 residue ubiquitinated in unloaded DmAgo1 by the E3 ligase Iruka^21^. Secondary structure designation is adopted from *Hs*Ago2 as represented in [5].

**Figure S2. The N-terminus of AtAGO1 is predicted to be an intrinsically disordered region**.

(A) Schematic representation of AtAGO1 with color-coding of the folded domains as indicated. Residues 1-268 were included in the IDR predictions in (B), (C), and (D).

(B) Intrinsic disorder prediction by IUPred^63^.

(C) Structure of the N-terminus predicted by Globplot^64^.

(D) Structure of the N-terminus predicted by PONDR^65^. The position of the N-coil is highlighted by the box.

**Figure S3. N-coil mutations reduce the strength of bimolecular fluorescence complementation mediated by AGO1-ATI1 and AGO1**^**Y691E**^**-ATI1**.

Half of the confocal microscopy images used for the quantification shown in Figure 5B have been assembled in this panel. KK, K185E/K190E.

**Figure S4**. ^**35**^**S-Met/Cys uptake in seedlings**.

Autoradiogram made from herbarium of *ago1-3*^-/-^/FLAG-*AGO1* seedlings 9 days after germination pulsed with ^35^S-Met/Cys for the indicated periods of time.

**Figure S5. Pulse labeling of AGO1**.

(A) Immunoblot analysis with *α*-FLAG antibody and autoradiogram of FLAG-AGO1, FLAG-AGO1^Y691E^ and FLAG-AGO1^KKME/Y691E^ pulsed with ^35^S-Met/Cys for the indicated periods of time. Amounts of FLAG-tagged AGO1 in total lysates and in immunoprecipitated fractions from seedlings using *α*-FLAG antibody are shown. * indicates a protein co-immunoprecipitating specifically with unloaded AGO1. The protein was identified as HSP70 as indicated by probing with *α*-HSP70 antibody. For this reason, ^35^S label from the ∼70 kDa band was not included in the AGO1 quantifications, see also Methods. KKME, K185E/K190E/M304E/E307A. Raw data from the gels in this panel underlie the graphical representations shown in Figure 6B,C.

(B) An independent repeat of the same type of experiment as in (C), with FLAG-*ago1*^Y691E^ and FLAG-*ago1*^KKME/Y691E^. Raw data from the gels in this panel underlie the graphical representation shown in Figure 6D.

**Table S1 Oligonucleotides used in the study**

