## Supplemental Figures for "Distinction between small RNA-bound and free ARGONAUTE via an N-terminal protein-protein interaction site"

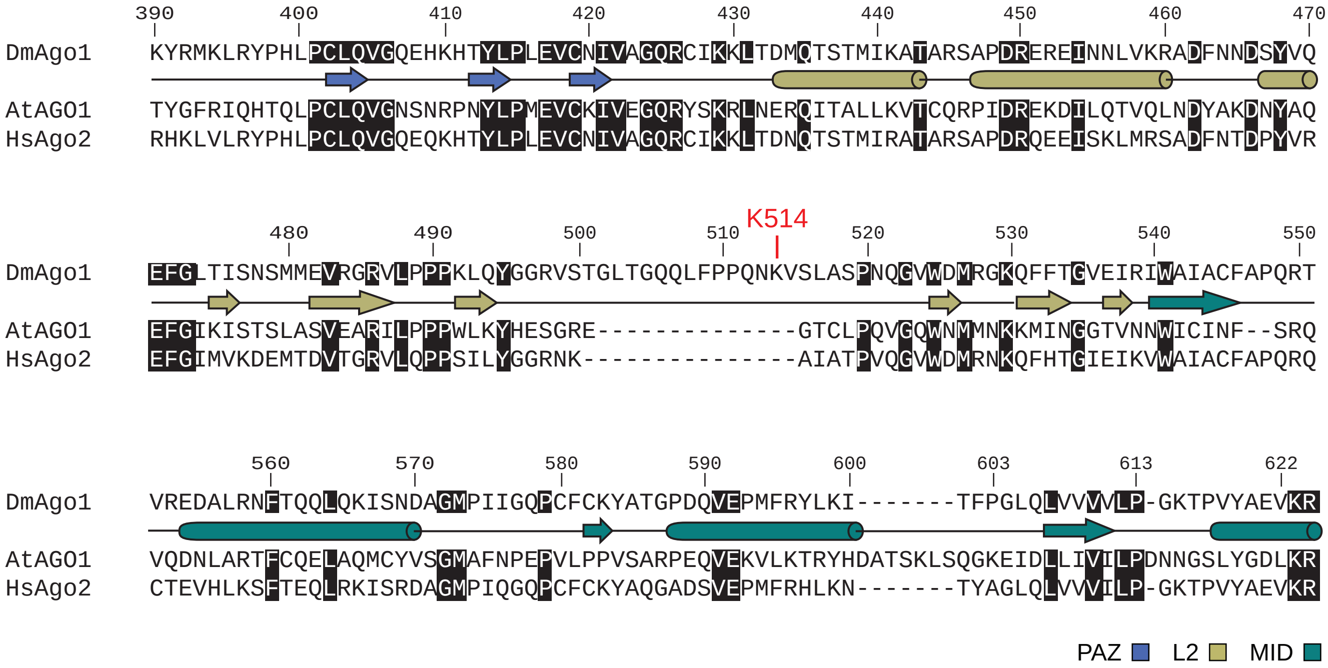


**Figure S1. Alignment of *Dm*Ago1 with *At*AGO1 and *Hs*Ago2.**
The alignment highlights the position of the Lys514 residue ubiquitinated in unloaded DmAgo1 by the E3 ligase Iruka[21]. Secondary structure designation is adopted from *Hs*Ago2 as represented in [5].

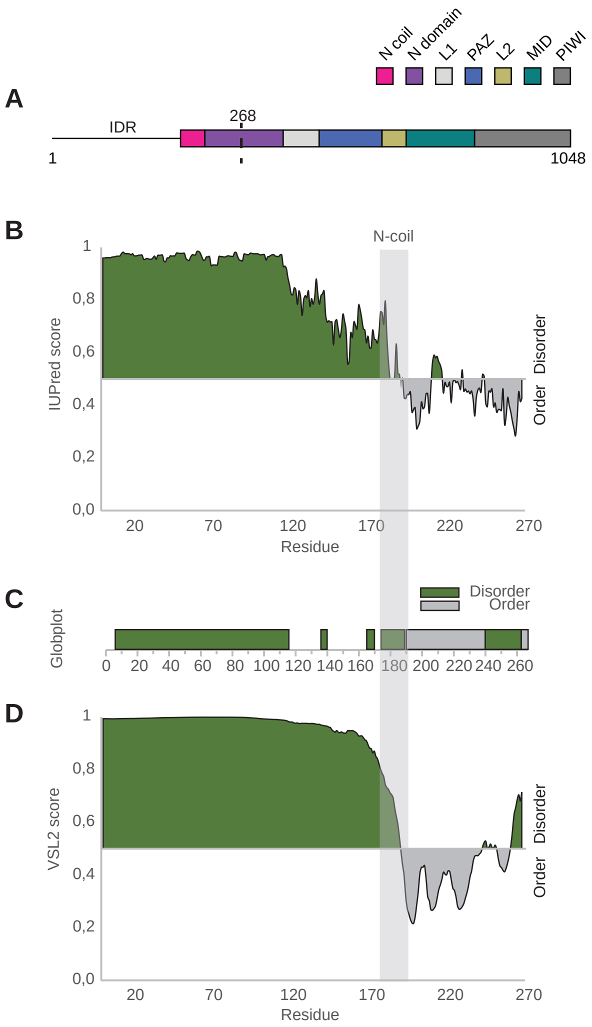


**Figure S2. The N-terminus of AtAGO1 is predicted to be an intrinsically disordered region.**
(A) Schematic representation of AtAGO1 with color-coding of the folded domains as indicated. Residues 1-268 were included in the IDR predictions in (B), (C), and (D).
(B) Intrinsic disorder prediction by IUPred[63].
(C) Structure of the N-terminus predicted by Globplot[64].
(D) Structure of the N-terminus predicted by PONDR[65]. The position of the N-coil is highlighted by the box.

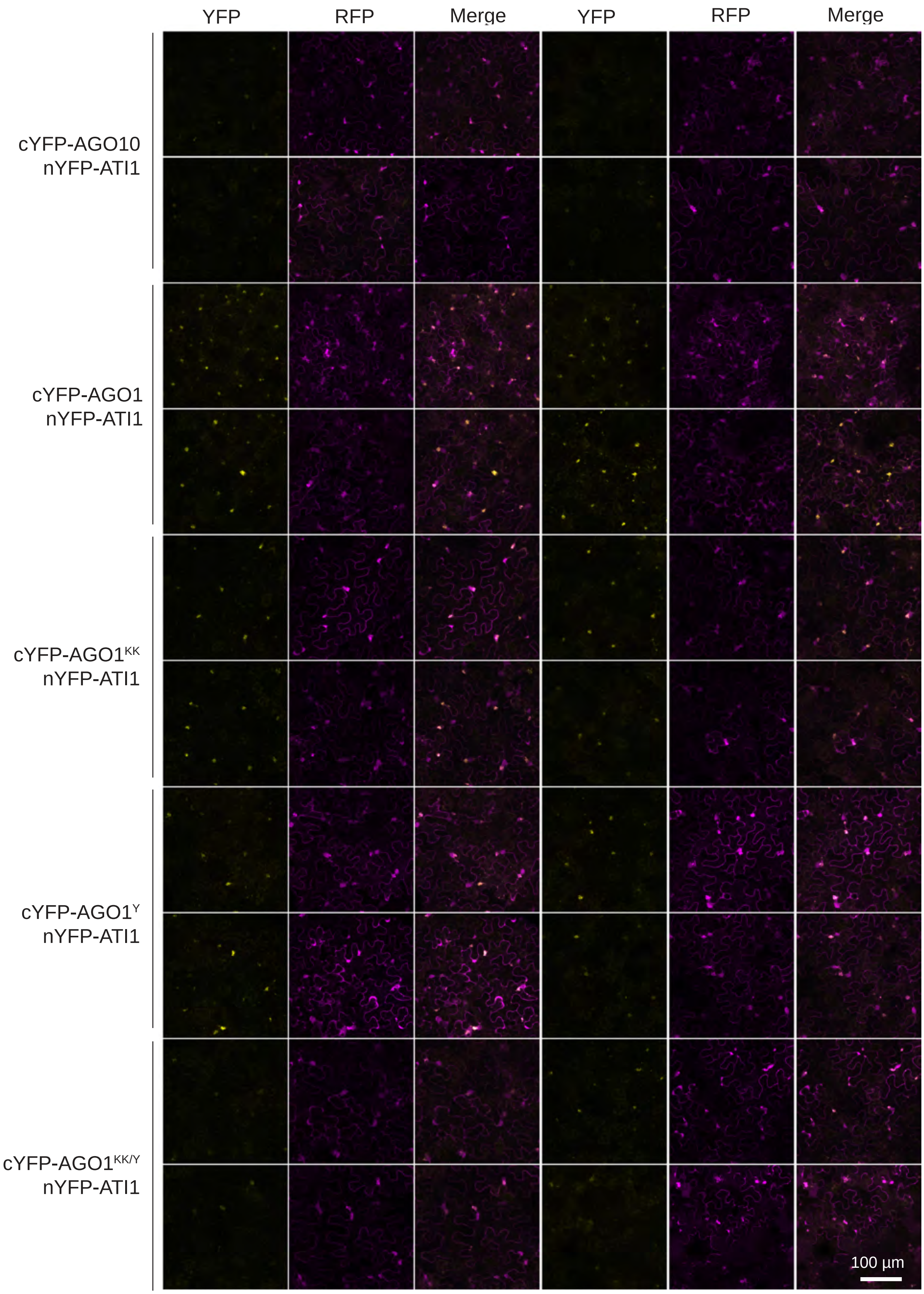


**Figure S3. N-coil mutations reduce the strength of bimolecular fluorescence complementation mediated by AGO1-ATI1 and AGO1$^{Y691E}$-ATI1.**
Half of the confocal microscopy images used for the quantification shown in Figure 5B have been assembled in this panel. KK, K185E/K190E.

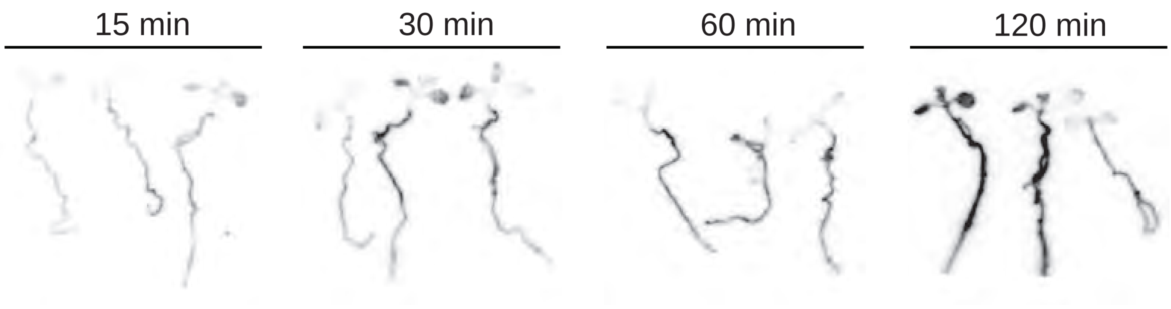


**Figure S4. $^{35}$S-Met/Cys uptake in seedlings.**
Autoradiogram made from herbarium of *ago1-3*$^{-/-}$/FLAG-*AGO1* seedlings 9 days after germination pulsed with $^{35}$S-Met/Cys for the indicated periods of time.

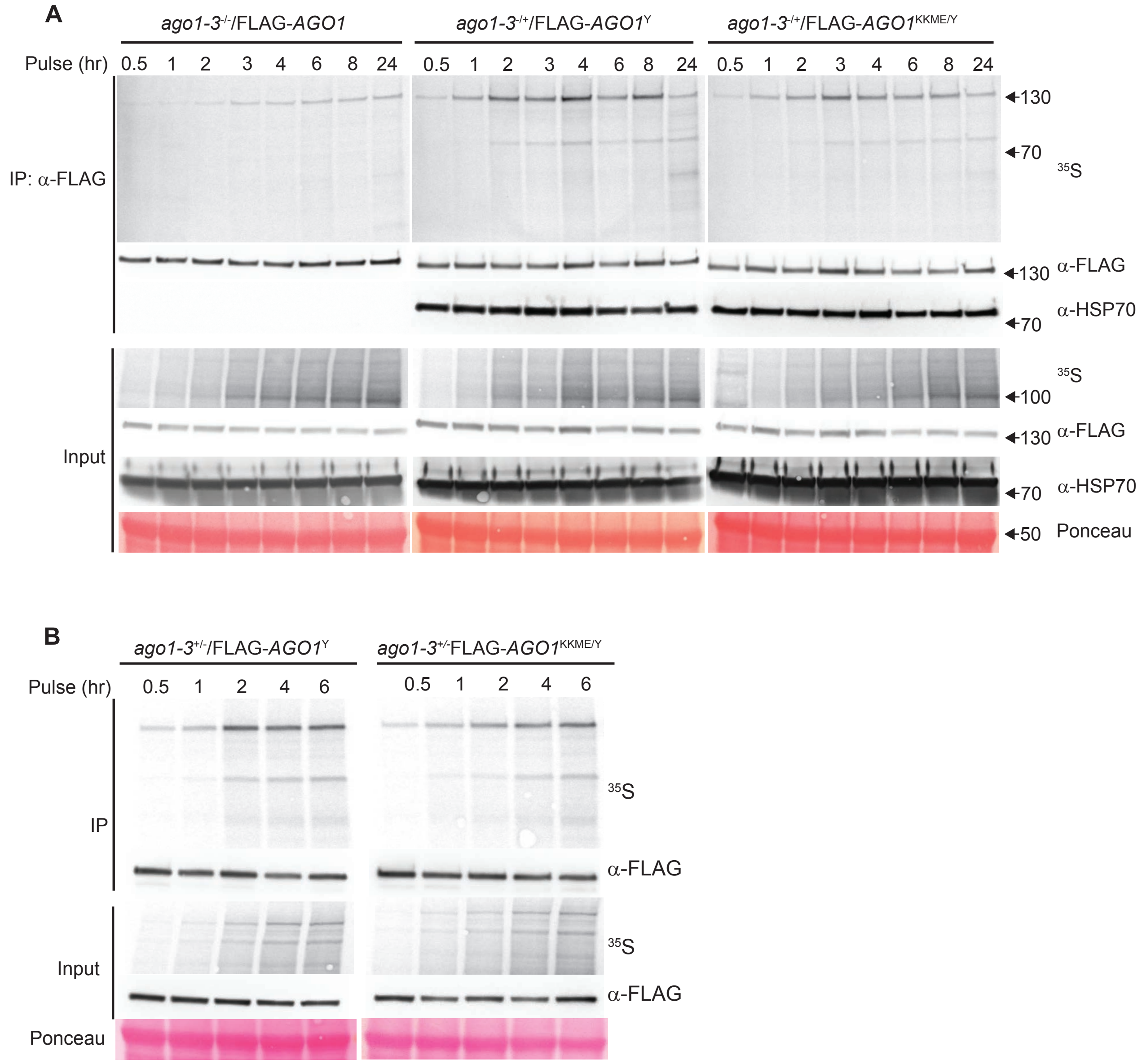


**Figure S5. Pulse labeling of AGO1.**

(A) Immunoblot analysis with α-FLAG antibody and autoradiogram of FLAG-AGO1, FLAG-AGO1$^{Y691E}$ and FLAG-AGO1$^{KKME/Y691E}$ pulsed with $^{35}$S-Met/Cys for the indicated periods of time. Amounts of FLAG-tagged AGO1 in total lysates and in immunoprecipitated fractions from seedlings using α-FLAG antibody are shown. * indicates a protein co-immunoprecipitating specifically with unloaded AGO1. The protein was identified as HSP70 as indicated by probing with α-HSP70 antibody. For this reason, $^{35}$S label from the ~70 kDa band was not included in the AGO1 quantifications, see also Methods. KKME, K185E/K190E/M304E/E307A. Raw data from the gels in this panel underlie the graphical representations shown in Figure 6B,C.
(B) An independent repeat of the same type of experiment as in (C), with FLAG-*ago1*$^{Y691E}$ and FLAG-*ago1*$^{KKME/Y691E}$. Raw data from the gels in this panel underlie the graphical representation shown in Figure 6D.
